# Determinism and stochasticity in seed dispersal-successional feedbacks

**DOI:** 10.1101/791988

**Authors:** Nohemi Huanca Nuñez, Robin L. Chazdon, Sabrina E. Russo

**Affiliations:** School of Biological Sciences, University of Nebraska, Lincoln, NE 68588-0118, USA; Department of Ecology and Evolutionary Biology, University of Connecticut, Storrs, Connecticut 06269-3043 USA

**Keywords:** Chronosequence, community assembly, Costa Rica, forest succession, secondary forest, seed rain, tropical forest

## Abstract

Regeneration of tropical secondary forests depends critically on seed input, and yet successional dynamics of seed dispersal remain poorly understood. We investigated the role of stochastic vs. deterministic processes in structuring seed rain in successional forests using four years of seed rain data collected at two time periods in four tropical secondary forest fragments representing a chronosequence and in mature forest. Determinism in successional trajectories is defined as predictable, directional, and orderly changes in community structure through time, resulting in convergence toward a climax community. We found that with increasing successional age, the community assembly of the seed rain in secondary forests became more deterministic, and community structure converged to that in the mature forest, both in terms of taxonomic and functional composition. Taxonomic similarity of the seed rain in successional forest to that of the mature forest increased with successional age, as did species co-occurrence and the percentage of shared species between the seed rain of successional and mature forests. The proportions of large, shade-tolerant species in the seed rain increased with successional age, although the proportion of animal-dispersed species increased only modestly. Analyses of the spatial variation in community structure in the seed rain among sites within each secondary forest showed evidence that assembly processes transitioned from being deterministic and convergent early on, to purely stochastic, and then to deterministic and divergent later in succession. Moreover, with increasing successional age, the composition of the seed rain became more similar to that of the mature woody stems in the forest, which could be an important deterministic driver of successional change, that, along with among site variation in landscape context and environment, could also generate idiosyncratic successional patterns among secondary forest fragments Our results suggest that the dominant processes influencing seed dispersal and assembly of the seed rain change during succession and point to successional feedbacks influencing the seed rain that are likely to shape regeneration trajectories.

## INTRODUCTION

Most of the original global extent of tropical forests (about 69%) has been lost to deforestation (FAO, 2018) caused by many processes, such as fires, hurricanes, and conversion to agricultural use (Brown and Lugo 1990; Guariguata and Ostertag 2001; Chazdon 2008). Despite the growing land area occupied by secondary forests, successional processes in tropical forests remain poorly known (Chazdon 2014; Norden *et al.*, 2015). Given the key role forests play in moderating local climate and ecosystem function (Hoffman *et al.*, 2003; Alkama and Cescatti 2016), as well as global carbon and other biogeochemical cycles (Ellison *et al.*, 2017), understanding how secondary forests regenerate is of critical importance.

Among the many ecological processes involved in forest regeneration (Foster and Tilman 2000; Guariguata and Ostertag 2001; Feldpausch *et al.*, 2005), seed input into deforested areas, particularly during early successional stages, can strongly influence successional trajectories (Svenning and Wright 2005). Seed dispersal is a critical process in forest regeneration because it determines colonization of deforested areas (Howe and Smallwood 1982; Levin *et al.*, 2003), establishes the initial spatial template for tree recruitment (Schupp *et al.* 2000; Russo and Augspurger 2004), and can affect the maintenance of species diversity (Hubbell 2001; Wright 2002; Terborgh *et al.*, 2017). Studies of tropical forest succession have focused on tree dynamics (Guariguata *et al.*, 1997; Chazdon 2008; Norden et al. 2017), with less attention focused on how seed dispersal affects succession (Costa *et al.*, 2012).

Seed dispersal is considered to be a highly stochastic process (Webb *et al.*, 2002; Lowe and McPeek, 2014), whereas there remains much disagreement as to whether succession is more deterministic or stochastic in space and time (Terborgh, *et al.*, 1996; Hubbell 2001). In the context of succession, deterministic processes are defined as predictable, directional, and orderly changes in species abundance and composition through time in a location that result in convergence toward a climax community, given a pool of potential colonizing species and a particular environmental setting (Clements 1916). On the other hand, stochastic processes are more strongly influenced by chance, and hence changes in species abundance and composition would be less predictable, with diverse behavior of individual species and less orderly successional trajectories that do not indicate convergence to a climax community (Gleason 1926). Seed rain into successional forests is influenced by many processes, including the proximity to and species composition of seed sources, the availability and diversity of seed dispersers, and the landscape context that influences movements of seed dispersers (Saunders *et al.*, 1991; Ricketts 2001). However, seed rain into secondary forests is also affected by the successional trajectory of the forest vegetation itself, creating a feedback between seed rain and forest successional processes that can not only affect forest regeneration, but also produce a deterministic successional signal through time in the seed rain.

The signature of deterministic and stochastic processes can also be seen spatially within a forest at a given time point. Deterministic patterns within a forest can take two basic forms: convergent and divergent. Convergent patterns, in which species composition across locations within a site is more similar than expected by chance, are assumed to be caused by abiotic filtering, which implicitly assumes that seeds are dispersed widely, and that filtering operates at the establishment and recruitment stages (Silvertown *et al.*, 2006; Webb *et al.*, 2006). However, in seed dispersal, a convergent pattern could result from any sort of filtering acting on the composition of the adult reproductive trees or limited access by seed dispersers that could cause greater compositional similarity of the seed rain within a site than expected. Divergent patterns, in which species composition among locations in a site are less similar than expected, are assumed to be the result of processes such as competitive exclusion (Kembel and Hubbell, 2006) and facilitative interactions (Valiente-Banuet and Verdú 2007). In seed dispersal, a divergent pattern could result from competitive exclusion of the adult reproductive trees or by the presence of a variety of seed dispersers. Since community assembly is influenced by both deterministic and stochastic processes (Adler *et al.*, 2007; Purves and Turnbull 2010), evaluating their relative importance in shaping the seed rain in secondary forests is key to understanding how seed dispersal affects vegetation recovery from disturbance.

In this study, we investigated the role of stochastic vs. deterministic processes in structuring seed rain in tropical successional forests during post-agricultural regeneration in Costa Rica. Using four years of data on seed rain of woody species collected at two time periods in forest fragments representing a temporal and spatial chronosequence with respect to successional age, we answered the following questions. First, how deterministic is the change in seed rain during tropical forest succession? If deterministic ecological processes predominate, then, assuming that a sufficiently wide range of successional ages of forests is captured, community structure of the seed rain in secondary forests with similar environments should converge towards that of the mature forest, regardless of initial differences. We would therefore expect the similarity of the seed rain between successional and mature forest to increase through time and between forests that are closer in successional age. Given that large seeds allow shade tolerance and are often animal-dispersed (Howe and Smallwood 1982; Westoby *et al.*, 1992; Grubb and Metcalfe 1996), we predicted that, as a proportion of the total seed rain, the quantity of seeds and species with large, animal dispersed seeds and shade-tolerant seedlings should increase with time and successional age. Second, does the relative importance of deterministic versus stochastic processes change during succession? We predicted assembly of the seed rain to be more stochastic in earlier than in later successional forests and to become more dominated by deterministic processes with time. Using a null model of seed rain, we tested whether observed dissimilarity decreased with time and successional age, indicating greater determinism. Given that a climax community is one that is self-replacing (Clements 1916; Denslow and Guzman 2000; Chazdon *et al.*, 2008), the species composition of the seed rain should become more similar with time and successional age to the composition of the woody species in the forest fragment, and the proportion of the seed rain comprised of immigrant species (species not locally represented at the site) should decline.

## METHODS

### Study site

This study was conducted in tropical, premontane, wet forest (Holdridge and Grenke, 1971) at La Selva Biological Station (hereafter, La Selva) and in surrounding areas in the Sarapiqui province of northeastern Costa Rica (Fig. S1). La Selva hosts a diversity of more than 1,850 plant species, with 350 species of trees. The dominant plants families are Pteridophyta, Orchidaceae, Araceae, Rubiaceae, Melastomataceae, Fabaceae, and Piperaceae, with *Welfia regia (Arecaceae), Socratea exorrhiza (Arecaceae)*, and *Pentaclethra macroloba* (Fabaceae) being the most abundant canopy species (Hartshorn and Himmel 1994). Mean annual rainfall and temperature is about ∼4000 mm and 26.5 C, respectively (Frankie et al. 1974). The areas surrounding La Selva once contained tropical forests with similar tree species composition, given the shared, regional tree species pool, but have experienced deforestation in the last several decades. Five 1-ha forest plots (50 m x 200 m) (Table 1; Fig. S1) were established in 1997 (four in secondary forest) and 2005 (one in mature forest) and have been censused annually for seedlings and trees (Menge and Chazdon, 2016). The plots are within a matrix of secondary, mature forest patches, agricultural areas (Norden *et al.*, 2012), and human settlements. The four secondary forest plots were used as cattle pastures after initial cutting of the mature forest and range in successional age (*i.e.*, time since the abandonment of pasturing) from 12 to 25 years old at the time of plot establishment (34 to 47 years old in 2019). The mature forest plot that has not been cleared or used for agriculture during modern times. Three plots are located inside La Selva, and two are ca. 6 km west of La Selva in privately owned farms.

**Table 1.** Stand characteristics of five 1-ha forest plots in successional and mature forest in Sarapiqui, Costa Rica. Plot names are those used in previous publications (Chazdon *et al* 2010). The number of trees per hectare is for stems ≥ 5 cm in diameter at breast height. La Selva BS indicates La Selva Biological Station.

| Plot abbreviation in 1997, 2017 | A1, A2 | B1, B2 | C1, C2 | D1, D2 | M |
| --- | --- | --- | --- | --- | --- |
| Successional age in 1997 (y) | 12 | 15 | 20 | 25 | Mature forest |
| Successional age in 2017 (y) | 32 | 35 | 40 | 45 | Mature forest |
| Plot Name | Lindero Sur | Tirimbina | El Peje secundario | Cuatro Rios | El Peje primario |
| No. trees/ha - 1997 | 1132 | 1071 | 1265 | 1140 | n.a |
| No. trees/ha - 2015 | 925 | 1020 | 1301 | 1256 | 1114 |
| No. woody species - 1997 | 64 | 100 | 110 | 136 | n.a |
| No. woody species - 2015 | 71 | 130 | 98 | 125 | 155 |
| Location | La Selva BS | Tirimbina | La Selva BS | Cuatro Rios | La Selva BS |
| Latitude, longitude | 10.41°N, -84.03°W | 10.40°N, -84.11°W | 10.43°N, -84.03°W | 10.39°N, -84.13°W | 10.42°N, -84.04°W |
| Surrounding landscape | Mature and secondary forest | Pasture, plantations, and secondary forest | Mature and secondary forest | Pasture, secondary and mature forest | Mature forest |

### Tree community and seed rain monitoring

All free-standing, woody stems in each forest plot with a diameter (dbh) ≥ 5 cm at breast height (1.3 m) have been tagged, mapped, and monitored for survival and growth annually since plot establishment in 1997 for secondary forest and 2005 for the mature forest (Chazdon *et al.*, 2005). The diameter of all living stems is measured to the nearest 0.1 mm, and new recruits (untagged trees with ≥ 5 dbh cm) are recorded at each forest plot.

Seed rain was quantified using the same methods for a total of 48 months over two time periods (24 months per time period). In each plot, ten 1-m^2^ seed traps (made from 1-mm fabric mesh suspended by a frame 1 m above the ground) were placed in a line down the middle of each plot every 20 m and 24.5 m from the plot edges. Traps were monitored monthly for 24 months from 1997 to 1999 in the four successional plots and for 24 months from 2015 to 2017 for the same four successional plots plus the mature forest plot. All contents of the traps were collected, and plant reproductive parts were separated from litter, sorted, and identified to the species level, and counted. Taxonomic names were standardized using the package ‘taxostand’ (Cayuela and Oksanen 2014) in R statistical software (R Core Development Team, 2019), and species were assigned codes for analyses (Table S1). Species with seeds ≤ 1 mm were excluded because seeds of this size could pass through the mesh and were not be reliably trapped. The length and width of a sample of seeds of each species was also measured (3-5 seeds per species). Seed length was aggregated into a categorical variable: small (≤ 6 mm) and large (> 6 mm) seeds (Grubb 1998; Saunders *et al.*, 1991). Based on literature and natural history (Santiago and Wright 2007; Comita *et al.* 2010; Sandor 2012; Wendt 2014), species in the seed rain were classified into two dispersal modes (animal-dispersed and non-animal dispersed) and two shade tolerance categories (light-demanding and shade tolerant).

### Statistical Analysis

All analyses were performed in R statistical software (R Core Development Team, 2019). We constructed community matrices of the seed rain abundance and incidence data and used these matrices for all analyses. Community matrices assembly is described in the supplementary methods section (Appendix S1). To evaluate how deterministic the change in seed rain is during tropical forest succession, we assessed the convergence of the community structure of the seed rain in successional forests towards that in the mature forest using three analyses. First, we evaluated how species composition of the seed rain based on abundance and incidence data varied across forests of different successional ages. We calculated multivariate distances in seed rain composition between each of the nine plots × time period (1997-1999, 2015-2017) combinations, using non-metric multidimensional scaling (NMDS) (Legendre and Legendre, 1998). Based on both presence/absence and abundance-weighted analyses, NMDS was implemented using the function *metaMDS* in the VEGAN package (v. 2.5-3) (Oksanen *et al.* 2018) with the Chao–Jaccard dissimilarity estimator (Chao *et al.*, 2005) and two dimensions. Results using other dissimilarity estimators were similar, so we only report those obtained with the Chao-Jaccard estimator. We visualized multivariate differences in the seed rain using biplots of NMDS components with 95% confidence ellipses based on the standard error, for each plot by time period combination. To test the statistical significance of differences in species composition, we used permutational multivariate analysis of variance (pMANOVA; Anderson, 2001), as implemented in the *adonis* function in the VEGAN package, with plot x time period combinations as factors. We used a post-hoc multilevel, pairwise analysis using the *pairwise.adonis* function (Martinez-Arbizu, 2017) to quantify differences between each plot within a time period and between time periods within a plot, correcting for multiple comparison with the Holm-Bonferroni correction.

To quantify the convergence of the successional forests to the mature forest, we constructed null models of species co-occurrences in the seed rain for the community matrix of each plot × time period combination. Null seed assemblages were constructed based on 1000 randomizations of the community matrix using the *randomizeMatrix* function and the independent swap algorithm (Gotelli 2000) in the package picante (v.1.8) (Kembel *et al.*, 2010), which randomizes the identities of the species in the seed rain in each community matrix while maintaining the same number of species and total number of seeds. Thus, only the species composition and abundances of the null seed rain assemblages could vary. The probability of co-occurrence in the seed rain of all possible pairs of species was calculated for each successional plot in each time period versus the mature forest using Schoener’s co-occurrence index (Pollock *et al.*, 2014) for the observed and null seed assemblages. Null seed assemblages were constructed only for the secondary forests, and the observed seed rain was used for the mature forest. Whether observed co-occurrence probabilities were different from those expected by chance was tested by comparison with the null distribution of co-occurrence indices based on a two-tailed test, using the 2.5th and 97.5th quantiles. The standardized effect size (SES) was calculated as the deviation of the observed value from the mean divided by the standard deviation from the random expectation, which allows comparison of co-occurrence values between sites and studies (Swenson 2014). A positive (negative) SES indicates a higher (lower) than the expected value of the co-occurrence index. We also calculated the percentages of species shared between each successional plot to the mature forest with estimates from the community matrices and the *vegdist* function with the Chao-Jaccard estimator in the VEGAN package. To compare species richness of functional groups based on seed size, shade tolerance, and dispersal mode represented in the seed rain across forests of different successional ages, we used rarefaction. Rarefaction curves were estimated using the iNEXT package in R (Chao and Jost 2012; Chao *et al.*, 2014) with 200 bootstrap replications. Additionally, for each forest plot, we extracted the rarefaction values by functional group and categories (*e.g.* seed size: large and small) and summed both categories to 100% to calculate the percentage of species by category, functional group, and forest plots. Then, we calculated a correlation between percentage of species in a specific category of the functional group by forest successional age.

If seed rain assembly converged deterministically, we expect the community structure of the seed rain in a plot should become more similar to that of mature forest as succession proceeds. First, this can be visualized in the NMDS biplot because multivariate distances are represented metrically (Brennan *et al.*, 2019). Second, the similarity index, the SES (standardized effect size value for each site; Swenson 2014) of co-occurrence index, and percentage of species shared between successional and mature forests should increase with time and successional age. Third, the similarity in the functional composition of successional with mature forest should increase through time, and there should be a greater compositional similarity between forests that are closer in successional age. Specifically, the number of seeds and richness of large, animal-dispersed seeds, and shade-tolerant species should increase with successional age.

To determine how the relative importance of deterministic and stochastic factors changed during succession, we used the following analyses. First, we assembled null seed rain communities using a modified version of the approach proposed by Chase *et al.*, (2011), which employs a modified Raup-Crick index (Raup and Crick 1979), taking into account species abundances (Stegen *et al.*, 2013; Alberti *et al.*, 2017). We conducted this null model analysis at two levels, within a forest for each pair of the ten traps and between pairs of forests traps. The null species composition was generated 9,999 times, and at each iteration, the abundance-weighted Bray-Curtis dissimilarity index between the two traps was calculated. This metric compares the measured β-diversity against the β-diversity that would be obtained under random community assembly by using a null distribution. The index was scaled to range from -1 to 1 by subtracting 0.5 and multiplying this difference by 2 (Chase *et al.*, 2011; Stegen *et al.*, 2013). Hereafter, we will refer to this metric as *RC*_*ab*_ (abundance-based Raup-Crick), which indicates whether a pair of traps is less similar (approaching 1), as similar (approaching 0), or more similar (approaching -1), than expected by chance. Values not different from zero indicate stochastic assembly of the seed rain, whereas values approaching 1 or -1 indicate predominance of processes causing divergence or convergence in seed rain composition, respectively. We regressed the mean *RC*_*ab*_ index of each pair of traps against successional age using a linear mixed model in the lme4 package with trap (or forest trap) as random effect. 95% confidence intervals on mean *RC*_*ab*_ values for forests of each successional age were calculated assuming a Student’s t-distribution.

Second, we quantified whether, during succession, the seed rain into a plot became increasingly dominated by the tree species present in the plot, indicating a seed rain-succession feedback involving a transition from seed rain sources outside the plot to sources inside the plot. Using ordinary least squares regression, we fitted a linear model of the species abundances in the seed rain as our dependent variable versus the tree community as our independent variable, using the tree community data for 1997-1998 and 2014-2015. Species not represented in the seed rain but present in the tree community, or vice versa, were assigned abundance of zero. We excluded lianas in this analysis because they were not included in plot censuses. We log-transformed the number of trees and number of seeds for values > 0, and zero abundance was left as zero. The slope of the relationship between the number of seeds of each species in the seed rain versus the number of adult individuals of each woody species was estimated for each plot by time period. We classified seeds with zero tree abundance as immigrant species, and the percentage of immigrants was regressed against forest successional age in a general linear model.

If the relative importance of deterministic processes increases during succession, we expect the following. First, the observed within-forest and among-forest *RC*_*ab*_ dissimilarity indexes should deviate increasingly from zero with successional age. Second, the correlation between species abundance in the seed rain and mature woody species in the forest should increase as succession proceeds. Third, the proportion of the seed rain that is comprised of immigrant species should decline with succession.

## RESULTS

Across all plots and time periods, the total seed rain was 53,552 seeds of 178 species (including 26 morpho-species), representing 51 angiosperm families (Table S1). Among successional forest plots, total seed accumulation in 24 months across all traps was generally higher in 1997 – 1999, than in 2015 – 2017 (Fig. S2a), with species such as *Casearia arborea* largely accounting for this difference. The average abundance of seeds after controlling for variation by species was significantly higher in 2015 – 2017 than in 1997 –1999 (*p* < 0.001) (Fig. S2b). Across all secondary forest plots and years, the two most abundant species in the seed rain were *Casearia arborea* and *Alchorneopsis floribunda* (Fig. S3), and four species were present in every month’s seed rain during the twelve months of 1997 – 1999 (*Aristolochia sprucei, Cordia bicolor, Pinzona coriacea*, and *Zanthoxylum sp.1*) and of 2015 – 2017 (*Euterpe precatoria, Goethalsia meiantha, Laetia procera* and *Welfia regia*) (Fig. S4).

### Convergence in seed rain composition over time during tropical forest succession

Species composition of the seed rain differed across forests of different successional ages in both time periods and between successional and mature forest. Based on both presence/absence and abundance-weighted analyses of all plot × time period combinations, the compositional differences revealed by community ordination were statistically significant (*F* = 8.3, *p* = 0.001 and *F* = 3.7, *p* = 0.001, respectively). Based on post-hoc analyses, the composition of the seed rain differed significantly between all pairs of plots × time period combinations, except between seed rain in the successional forests C and D in 2015 – 2017, which were only marginally different (Table S2). Visual inspection of community ordination showed that the species composition of seed rain in successional forests largely converged towards that in the mature forest after 20 years of succession (Fig. 1). Successional forests sampled in 1997 – 1999 (A1-D1 in red) clustered together in a distinct region of the ordination space compared to the same forests sampled in 2015 – 2017 (A2-D2 in blue), which were closer in ordination space to the mature forest (M in black) (Fig.1). Shade tolerant species (green font) were present in the seed rain in both 1997 – 1999 and 2015 – 2017 but were more prevalent in successional forests in 2015 – 2017. Light demanding species (purple font) were abundant across both 1997 – 1999 and 2015 – 2017 but were prevalent in the successional forests in 1997 – 1999.

**Figure 1.**
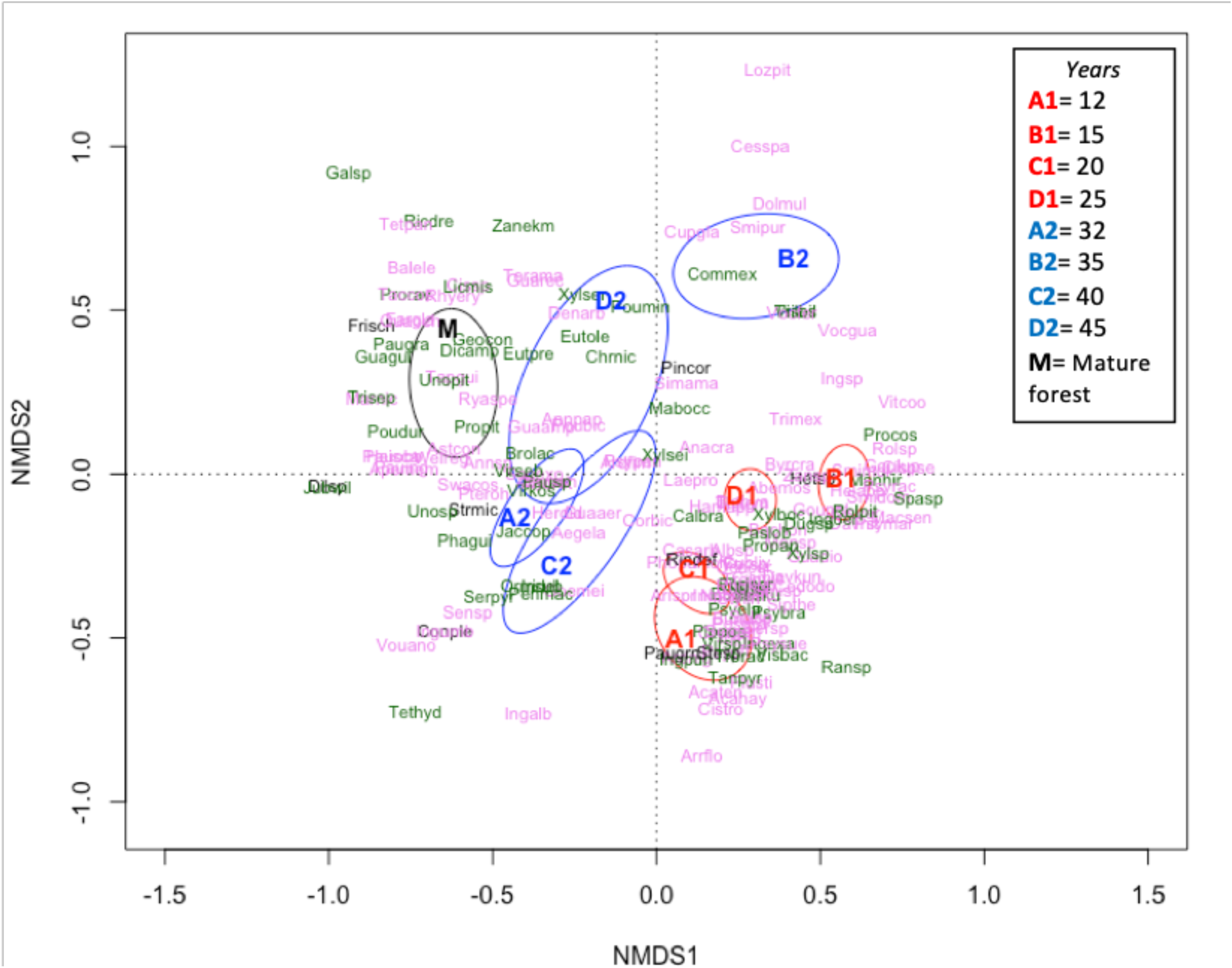
Differences in species composition of the seed rain across four successional and mature forest plots over two time periods in Sarapiquí, Costa Rica. Successional plots were sampled in 1997-1999 (A1-D1 in red) and 2015-2017 (A2-D2 in blue), and the mature forest (M in black) was sampled only in 2015-2017. Species abbreviations correspond to the first three letters of the genus and species, given in Table S1, with green font indicating shade tolerant species and purple font indicating light demanding species. Ellipses are 95% confidence ellipses based on standard errors. Plots A, C, and M are located inside LSBS.

The standardized effect size (SES) of the co-occurrence of species in the seed rain of successional and mature forests, was significantly lower than expected based on random assembly (Fig. S5), indicating that seed rain in successional forests represented a subset of the species found in the seed assemblages at the mature forests. Moreover, the negative SES declined significantly with successional age (*R* ^*2*^ = 0.67, *p* < 0.01) (Fig. 2a), indicating greater species co-occurrence in the seed rain of successional and mature forests. After 20 years of succession, the percentage of species shared in the seed rain between successional forests and the mature forest increased (*t* = 3.9, *p* = 0.03) (Fig. 2b, Table S3).

**Figure 2.**
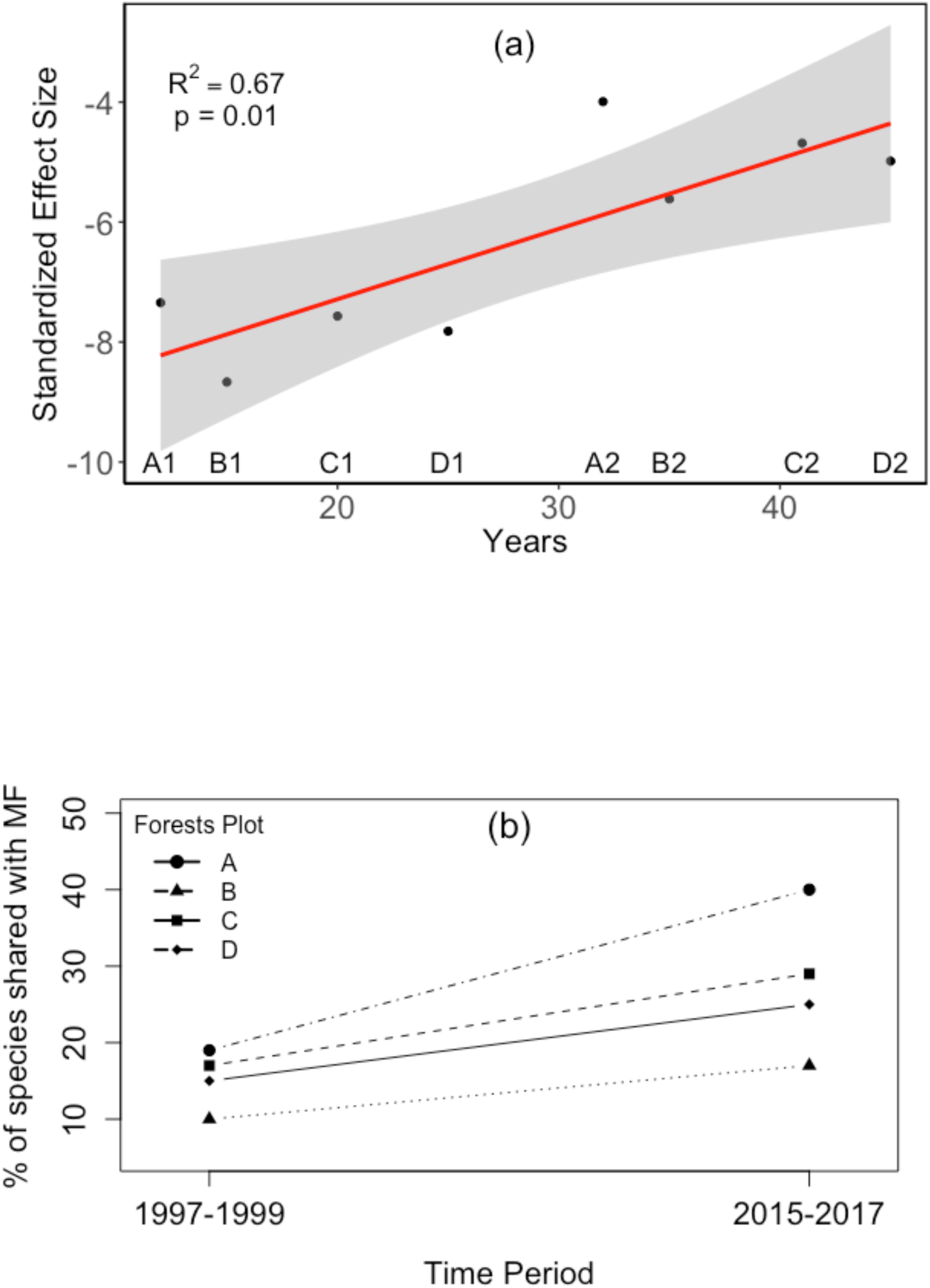
Convergence of species composition of the seed rain in secondary forests to that of the mature tropical forest during succession in Sarapiquí, Costa Rica. (a) Changes in standard effect size (SES) during succession for co-occurrence of species in the seed rain of successional and mature forests. A1, B1, C1, D1 and A2, B2, C2, D2 represent the 1997-1999 and 2015-2017 time periods, respectively. More negative values of SES indicate lower-than-average expected co-occurrence with the mature forest. The red line is the ordinary least squares regression line with the 95% confidence interval on the model fit indicated in gray shading. (b) Percentage of species in the seed rain shared by each successional forest with the mature forest in 1997-1999 and 2015-2017.

Rarefaction curves of species by functional groups showed successional forests converging towards the composition of the mature forest with time (Fig. S6). In successional forests, species richness of small-seeded (≤ 6 mm), light-demanding and non-animal dispersed was higher in the seed rain in 1997 – 1999 than in 2015 – 2017 and mature forests (Figs. S6a, S6c and S6e). Whereas the reverse was true for the richness of large-seeded and shade-tolerant species, when comparing each forest plot in 1997 – 1999 and 2015 – 2017, with mature forests having the highest richness (Figs. S6b and S6d). The patterns in the animal-dispersed species were more variable (Figs. S6e and S6f). The percentage of species in the seed rain within each forest plot of large and shade tolerant species increased with successional age, whereas the percentages of small and light demanding species decreased (Fig. 3), although for shade tolerance, the change was only marginally significant. Meanwhile, the trend for animal and non-animal dispersed was not statistically significant.

**Figure 3.**
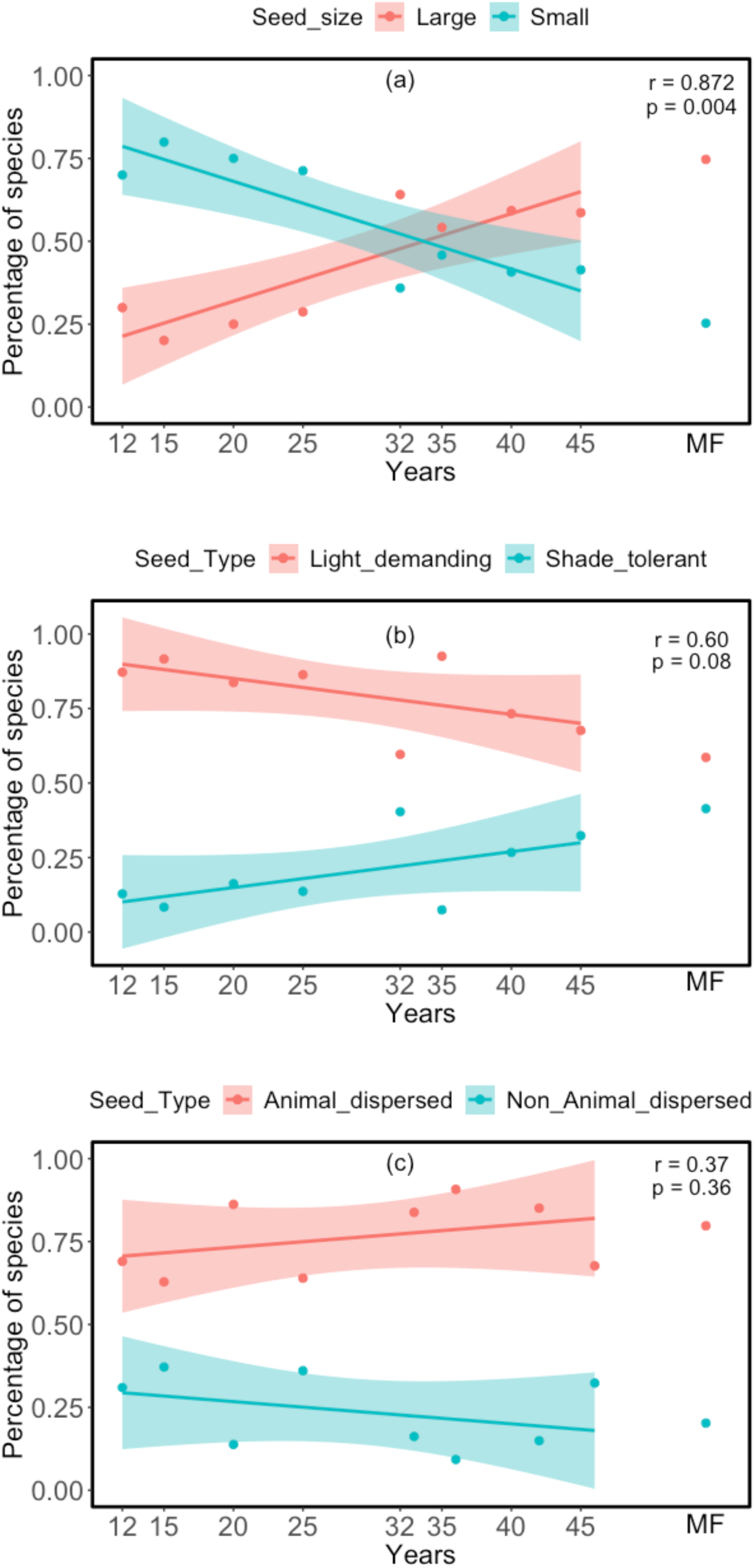
Changes in the functional composition of the seed rain in secondary forests of increasing successional age in Sarapiqui, Costa Rica. Shifts in the percentages of: (a) species with small (≤ 6 mm) and large seeds (> 6 mm); (b) shade tolerant and light demanding species; (c) animal-dispersed and non-animal dispersed species represented in the seed rain. (a) small (≤ 6 mm) and large seeds (> 6 mm) within each forest plot; (b) seeds of shade tolerant and light demanding species; (c) seeds of animal-dispersed and non-animal dispersed species. The percentage of species by category is based on calculated asymptotic Chao diversity estimates from rarefactions (Fig. S6), converted to percentages. Since there are only two categories, the correlation was conducted for the category that was increasing.

### Temporal change in determinism versus stochasticity of the seed rain during tropical forest succession

There was a positive, statistically significant relationship between the abundance-based Raup-Crick (*RC*_*ab*_), measuring within-site variation in spatial community structure of the seed rain, and forest successional age (*R*^*2*^ = 0.33, *p* < 0.01) (Fig. 4a), from values close to -1 for younger forests to values close to 1 for older forests. The *RC*_*ab*_ for the seed rain of the 12-year-old forest was significantly less than zero, indicating a more similar community structure of the seed rain than expected by chance (*i.e.*, low variability in the seed rain structure across traps within this forest). However, the *RC*_*ab*_ values for the seed rain in the 20, 25, 33, and 36-year-old forests did not differ from zero, consistent with stochastic assembly. In contrast, the *RC*_*ab*_ values for the seed rain in the 42- and 46-year-old forests were greater than zero, indicating a less similar community structure of the seed rain than expected by chance (*i.e.*, high variability in the seed rain structure across traps within the forest). Comparing each plot after 20 years of succession, there was no relationship between the *RC*_*ab*_ measuring temporal turnover and forest successional age (Fig. 4b). However, all *RC*_*ab*_ indexes measuring temporal turnover were greater than zero, indicating that over time each plot’s similarity to itself declined more than expected by chance.

**Figure 4.**
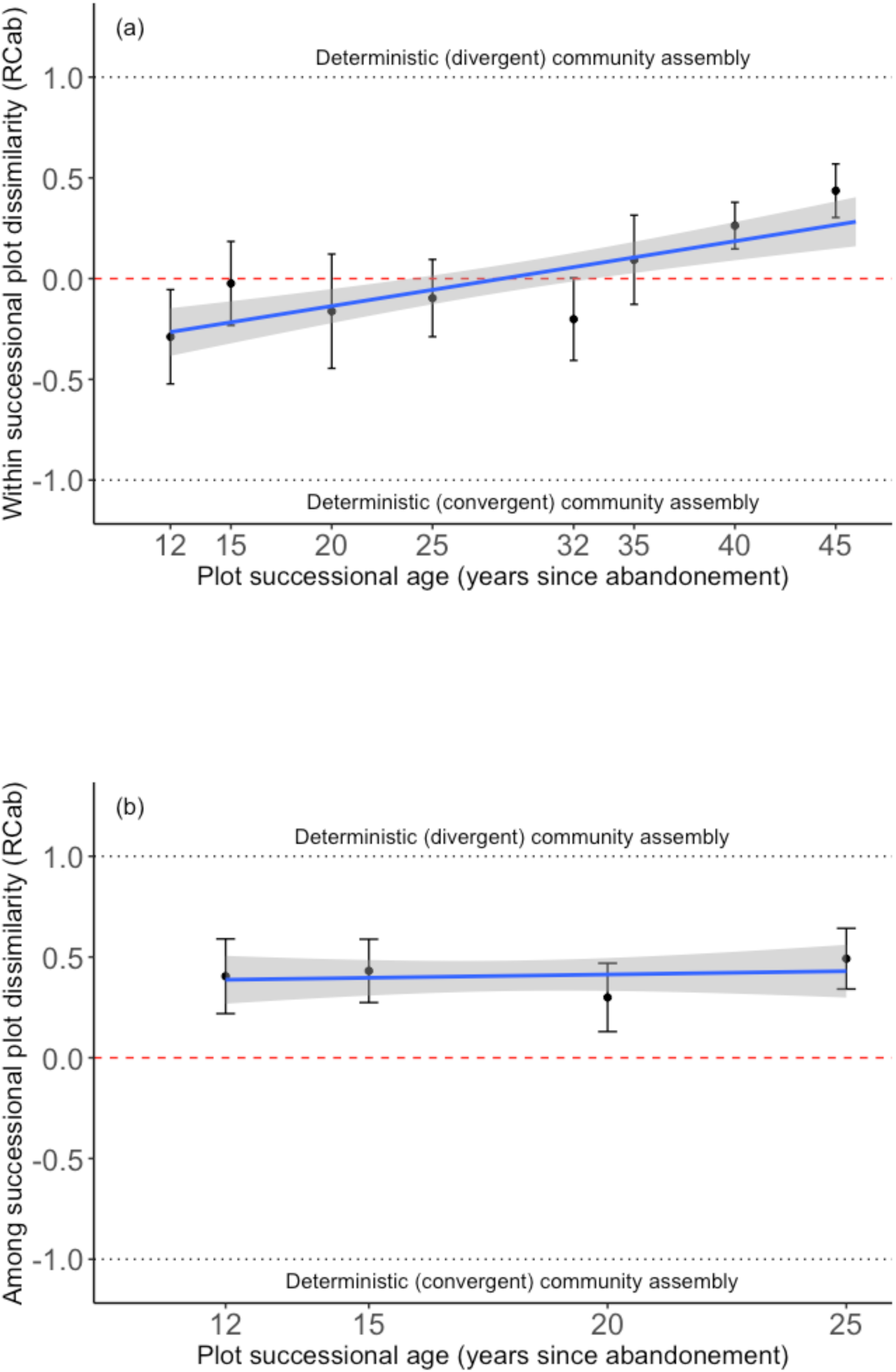
Increasing determinism of the seed rain during succession based on the abundance-weighted Raup-Crick dissimilarity index (*RC*_*ab*_). *RC*_*ab*_ values range from -1 to 1, indicating whether community structure within plots across all successional ages (a) or between time periods for the same plot (turnover, (b)) are more dissimilar (approaching 1), as dissimilar (approaching 0), or more similar (approaching -1), than expected by chance. The red horizontal line at *RC*_*ab*_ of zero denotes purely stochastic community assembly. Points are mean trap *RC*_*ab*_ values, error bars are the 95% confidence interval. (a) Variation of within-plot spatial community structure of the seed rain with successional age of the forest. (b) Temporal turnover (between time periods) of *RC*_*ab*_ with successional age. Forests were sampled in two time periods, 1997-1999 and 2015-2017, and *RC*_*ab*_ was estimated between the two time periods for each forest plot.

The seed rain in the mature forest had a strong relationship to its own tree composition, but the relationships varied for the successional forests and between the two time periods (Fig. 5). In 1997 – 1999, species abundances in the seed rain were not significantly related to those in the tree community (Fig. 5, panel A1, B1, C1, D1). However, after 20 years of succession, species abundance in the tree community became a better predictor of species abundance in the seed rain. Estimates of the slope of this relationship were significantly positive for all successional forests in 2015 – 2017, with the exception of forest B, which was only marginal (Fig. 5, panel B2). The proportion of immigrant seeds decreased with successional age (*R* ^*2*^ = 0.78, *p* < 0.01) (Fig. S7).

**Figure 5.**
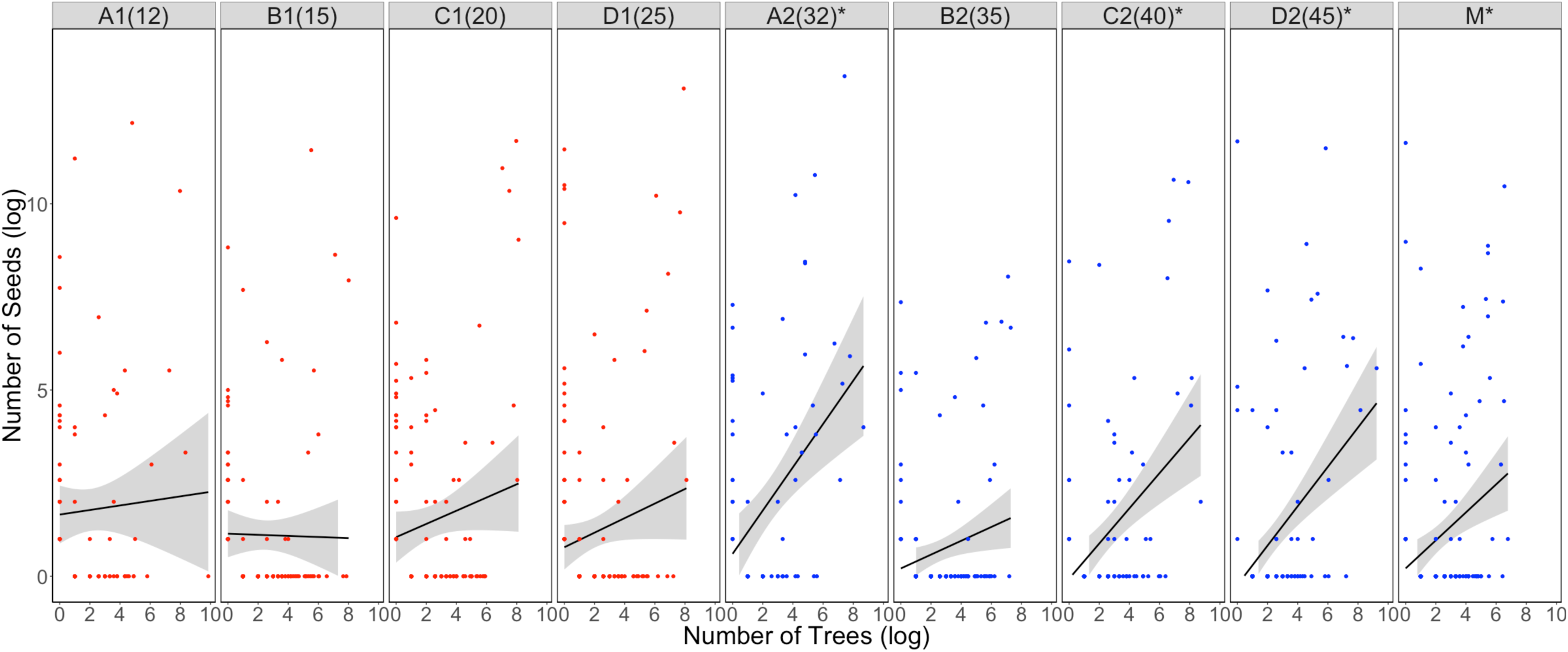
Positive seed rain-succession feedback increasing with successional age of the forest, as indicated by the relationship between the number of stems and the number of seeds of each woody species. Different panels correspond to forests of different successional ages. Non-zero values were log-transformed, so that a value of 0 is equivalent to absence of that species in either the seed rain or tree community. The asterisk indicates that the slope parameter was significantly different from zero (2015-2017: A2, *p <* 0.01, slope = 0.58; B2, *p =* 0.08, slope = 0.19; C2, *p* < 0.01, slope = 0.47; D2, *p* < 0.01, slope = 0.53; E, *p*<0.01, slope= 0.38). Successional age is in parentheses after the abbreviation, except for the mature forest (M) (Table 1).

## DISCUSSION

Reforestation after abandonment of former agricultural land depends critically on the input of tree seeds, yet how forest successional processes feedback to structure the quantity, as well as taxonomic and functional diversity, of seed rain during regeneration is not well understood. We sampled the seed rain into tropical secondary forests of varying successional ages in a chronosequence at two time periods and found that while seed rain retained some signatures of stochasticity, it became more deterministic with increasing successional age of the forest. Although these secondary forests differ in proximity to seed sources and the species composition of those sources, the taxonomic composition of the seed rain into them showed convergence toward that of the mature forest as succession progressed. This increase in determinism with succession was also manifested in terms of functional diversity, as significant shifts in seed size and ecological strategy to greater proportions of larger-seeded and shade-tolerant species, but only non-significant increases in the proportions of animal species vs non-animal dispersed species. The taxonomic and functional composition of the seed rain largely converged with that of the mature forest, although individual forests exhibited idiosyncratic trends, likely owing to diverse factors related to landscape context and site history. While the community structure of the seed rain within each secondary forest was spatially convergent at earlier successional ages, at later ages, it became more spatially divergent, consistent with the complex spatial structuring of diversity characteristic of mature tropical forests. Nonetheless, the composition of the seed rain became more similar to that of the mature trees in the forest with increasing successional age. Our results imply that an important driver of increasing determinism of the seed rain during succession is seed-rain successional feedbacks, which are likely to shape regeneration trajectories, particularly in secondary tropical forests.

### Successional convergence in species composition of the seed rain

Previous succession studies using a time-series approach have not always found convergence in species composition to that of the mature forest that is often observed in chronosequence studies (Peña-Claros 2003; Chazdon *et al.*, 2003; Lebrija-Trejos *et al.*, 2010). However, our results comparing similarity in community structure of the seed rain in successional forests showed a clearer pattern of convergence towards that of the mature forest in the temporal scale (1997 – 1999 to 2015 – 2017) than in the chronosequence scale (A1 to D1 and A2 to D2), indicating that successional age is not the only factor leading to differences between sites at any point in time. This result was supported by visual inspection of the metric ordination of the seed rain, in which seed rain composition in 2015 – 2017 was more similar to mature forest than it was in 1997 – 1999 in every successional forest. This was particularly true of plots A and C, which like the mature forest (M), are inside La Selva Biological Station. In contrast, while seed rain composition of plots B and D, which are outside of La Selva Biological Station, moved towards that of the mature forest, the shifts were not as direct as for plots A and C. The co-occurrence of and percentage of shared species in the seed rain of mature and successional forests increased with successional age, indicating gradual convergence towards the community structure of the mature forest. However, seed rain in successional forests still shared few species with the mature forest than expected, suggesting that even 45-year-old secondary forests still require substantial time to reach a mature or climax state. As with the ordination, this pattern was more strongly observed comparing successional forests through time, as co-occurrence and the percentage of shared species increased consistently for every successional forest from 1997 – 1999 to 2015 – 2017, in contrast to comparisons at the chronosequence scale. Thus, our results show that seed communities became more similar to the mature forest over time during forest regeneration, as successional theory predicts, but that there were also idiosyncratic patterns among secondary forests. Variation in the successional trajectories of different forests may arise from site-specific differences in landscape context, such as proximity to seed sources, the species composition of seed sources, visitation by dispersal agents, and the surrounding matrix (Ricketts, 2001), as well as in varying contributions of rare versus common and generalists versus specialist woody species to stochasticity or determinism (Norden *et al.* 2017). We sampled only one mature forest site in the area near the successional forests, but there are environment-driven differences in composition of the mature forest, even within a few kilometers distance (*e.g.*, Clark *et al*., 1999). The inclusion of more than one mature forest would have clarified convergence patterns more accurately.

### Successional convergence in functional composition of the seed rain

Trait-based analyses of successional trajectories have often shown that functional properties can exhibit more predictable directional changes indicating convergence to mature forest, compared to taxon-based analyses (Chazdon 2008; Norden *et al.*, 2012; Chazdon 2014, but see Boukili and Chazdon 2017). Our results generally confirmed these findings in that we found the functional composition of the seed rain in successional forests to converge towards that of the mature forest with time. The percentages of both large-seeded and shade tolerant species increased from 1997 – 1999 to 2015 – 2017. The richness of light-demanding species was higher in the seed rain in successional forests in 1997 – 1999, compared to 2015 – 2017 and in the successional forests in both years, compared with the mature forest. These results reflect successional feedbacks, as shorter-lived, light-demanding species (Bazzaz and Pickett 1980; Wright *et al.*, 2010) are replaced by longer-lived, more shade-tolerant species. Succession therefore increasingly favors the recruitment and maturation of, and hence, seed input, by, larger seeded species, as understory light levels decline, and the canopy closes, reducing the representation of small-seeded, light-demanding species among the mature trees contributing seed rain in the forest. Functional convergence in terms of dispersal modes represented in the seed rain was less distinct, consistent with the facts that there are many non-animal dispersed species in mature forests and that many animal seed dispersers are known to visit successional, as well as mature, forests. For example, phyllostomid bats are thought to make important contributions to seed rain in early successional forests because they consume fruits and disperse seeds of early successional species and fly across forest fragments (Muscarella and Fleming, 2007). Small, tent-roosting bats, such as *Artibeus watsoni*, promote dispersal of larger seeded species abundant mostly in late successional forest (Melo *et al.*, 2009). Thus, due to variable specialization of some animal species to successional versus mature forests, functional composition of the seed rain in terms of dispersal mode may not converge reliably to that of mature forest.

### Stochasticity and determinism in the seed rain: changes with forest successional age

We found that stochasticity and determinism in the spatial patterns in the community structure of the seed rain within each forest changed with successional age. In the youngest forests, community structure of the seed rain was deterministic, but spatially convergent, which could be caused by environmental constraints on community composition of woody species or the filtering out of species from the regional pool that could not arrive due to dispersal limitation. In middle-aged forests, community structure of the seed rain was more stochastic. In contrast, in the oldest successional forests, it was deterministic, but spatially divergent, which may occur when biotic interactions favor niche differentiated species. Consistent with these possible explanations, Norden *et al.* 2012 found that tree species with high mortality rates during early stages of tropical forest succession were more closely related than tree species that survive, and tree species that join the community were more distantly related than persisting species, indicating increasing divergence of the composition of the woody stems in the forest as succession proceeds, consistent with our findings for seed rain. We also observed analogous patterns in the temporal turnover within each forest. Namely, after 20 years of succession (2015 – 2017) there was greater divergence in community structure of the seed rain within forests than in 1997 – 1999, even though the number of species in the seed rain within each secondary forest plot in the 2015 – 2017 was not higher than in 1997 – 1999. Because biotic interactions change during succession (Chazdon 2008), the increasing importance of deterministic, compared to stochastic, processes during succession could be the result of a shift from interactions with more generalist, to more specialist, associations (Büchi and Vuilleumier 2014), which may lead to more deterministic outcomes of interactions with antagonists (e.g., competitors, parasites, herbivores, pests, and pathogens) and mutualists (e.g., frugivores, mycorrhizal fungi). Lastly, the species composition of the seed rain became increasingly more similar through succession to the composition of the adult trees in the plant community. So, the proportion of the seed rain that is comprised of immigrant species declined with succession, which indicates that seeds in late successional and mature forests are composed principally of seeds deposited by parent trees growing locally within the forest, consistent with the interpretation that specialist species of mature forest are dispersal limited (Norden *et al.* 2017). Since the woody stems themselves are also becoming more similar in composition to the mature forest (Letcher and Chazdon 2009), our results are indicative of a feedback between seed dispersal and forest successional processes.

### Conclusion

Seed dispersal is a process that is often considered highly stochastic. Even so, comparing the community structure of the seed rain in secondary forests using combined data from chronosequences and time series revealed significant determinism in seed rain that increased as succession progressed in these Costa Rican forests. At least two mechanisms may account for this. First, seed rain-successional feedbacks may cause the seed rain to become ever-more strongly determined by the local tree community, rather than immigrant seeds, feedbacks that may be particularly influential in secondary tropical forests. Second, changes in biotic interactions from generalist to specialist may cause the outcomes to become more deterministic. Here, we have shown evidence for the first mechanism. Future studies should examine whether seed dispersal interaction networks change from greater connectivity earlier to greater modularity or nestedness later in succession, which would provide evidence of more frequent specialist interactions that could lead to the increasing determinism of the seed rain during succession that we observed.

## ACKNOWLEDGEMENTS

We thank J. Paniagua, B. Paniagua, and E. Salicetti for field assistance. Seed rain data from 1997-1999 were collected by T. Robles-Cordero with assistance from J. Paniagua, M. Molina, and J. Romero. We thank the Organization for Tropical Studies (OTS) and local landowners for permission to conduct the research. The OTS David and Deborah Clark Fund, University of Nebraska – Lincoln, Walker Fellowship for Graduate Students in Botany and the National Geographic Society funded this research. Censuses in the forest plots were funded by the Andrew W. Mellon Foundation, NSF DEB-0424767, NSF DEB-0639393, NSF DEB-1147429, NASA Terrestrial Ecology Program, University of Connecticut Research Foundation, National Geographic Society, and Costa Rica Debt for Nature Program. We thank J. Stegen for sharing programming code.

