## Supplemental material 1 for "Determinism and stochasticity in seed dispersal-successional feedbacks"

**Appendix S1: Supplemental Methods Section**

***Statistical Analysis: Constructing seed rain community matrices***

All analyses were performed in R statistical software (R Core Development Team, 2019). We constructed community matrices of the seed rain data in which each cell is the total number of all seeds of each species recorded in each trap in a plot, summed over the 24 monitoring months in a time period, separately for time periods 1997-1999 and 2015-2017 for a total of nine abundance community matrices. Based on the abundance matrices, we created incidence matrices (presence/absence of species within a trap). We used these matrices for all analyses. Although the seed rain in the mature forest was only quantified for time period 2015-2017, we assumed that the composition of the seed rain of the mature forest would be similar in 1997-1999 since no major disturbances or changes in forest structure have occurred to the mature forest plot. We therefore compared data from the successional plots in both time periods with the mature forest data collected in 2015-2017.

Using only successional plots at each time period (1997-1999 and 2015-2017), we determined species present during the 12 consecutive months at each time period. Also, using a generalized negative binomial model, where number of seeds was the response variable, trap and time periods were fixed factors and plot, month and species as random factors, we calculated which time period had higher species seed abundance by $m^2$. A similar model where plot and time period were fixed factors and, trap, month and species were random effects, calculated the species with highest and lowest abundance across all the successional plot.

**Table S1.** List of codes, scientific names, and family for all species in this study.

| N | Code | Genus | Species | Family |
|---|---|---|---|---|
| 1 | Abemos | *Abelmoschus* | *moschatus* | Malvaceae |
| 2 | Abupan | *Abuta* | *panamensis* | Menispermaceae |
| 3 | Acahay | *Acacia* | *hayesii* | Leguminosae |
| 4 | Acaten | *Acacia* | *tenuifolia* | Leguminosae |
| 5 | Aegela | *Aegiphila* | *elata* | Lamiaceae |
| 6 | Albsp | *Albizia* | *sp* | Leguminosae |
| 7 | Alcflo | *Alchorneopsis* | *floribunda* | Euphorbiaceae |
| 8 | Allplu | *Allomarkgrafia* | *plumeriiflora* | Apocynaceae |
| 9 | Anacra | *Anaxagorea* | *crassipetala* | Annonaceae |
| 10 | Annpap | *Annona* | *papilionella* | Annonaceae |
| 11 | Annsp | *Annona* | *sp* | Annonaceae |
| 12 | Anoret | *Anomospermum* | *reticulatum* | Menispermaceae |
| 13 | Apemem | *Apeiba* | *membranacea* | Malvaceae |
| 14 | Ardfim | *Ardisia* | *fimbrillifera* | Primulaceae |
| 15 | Ardnig | *Ardisia* | *nigropunctata* | Primulaceae |
| 16 | Arispr | *Aristolochia* | *sprucei* | Aristolochiaceae |
| 17 | Arrflo | *Arrabidea* | *florida* | Bignoniaceae |
| 18 | Astcon | *Astrocaryum* | *confertum* | Arecaceae |
| 19 | Bachon | *Bactris* | *hondurensis* | Arecaceae |
| 20 | Balele | *Balizia* | *elegans* | Leguminosae |
| 21 | Bighya | *Bignonia* | *hyacinthina* | Bignoniaceae |
| 22 | Brolac | *Brosimum* | *lactescens* | Moraceae |
| 23 | Bunoce | *Bunchonsia* | *ocellata* | Malpighiaceae |
| 24 | Byrart | *Byrsonima* | *arthropoda* | Malpighiaceae |
| 25 | Byrcra | *Byrsonima* | *crassifolia* | Malpighiaceae |
| 26 | Calbra | *Calophyllum* | *brasiliense* | Clusiaceae |
| 27 | Casarb | *Casearia* | *arborea* | Salicaceae |
| 28 | Cedodo | *Cedrela* | *odorata* | Meliaceae |
| 29 | Cesspa | *Cespedesia* | *spathulata* | Ochnaceae |
| 30 | Chrnic | *Chrysochlamys* | *nicaraguensis* | Clusiaceae |
| 31 | Cismic | *Cissus* | *microcarpa* | Vitaceae |
| 32 | Cispse | *Cissus* | *pseudocyoides* | Vitaceae |
| 33 | Cissp | *Cissus* | *sp* | Vitaceae |
| 34 | Cistro | *Cissampelos* | *tropaeolifolia* | Menispermaceae |
| 35 | Cisver | *Cissus* | *verticillata* | Vitaceae |
| 36 | Commex | *Compsoneura* | *mexicana* | Myristicaceae |
| 37 | Conple | *Conceveiba* | *pleiostemona* | Euphorbiaceae |
| 38 | Corbic | *Cordia* | *bicolor* | Boraginaceae |
| 39 | Crywar | *Cryosophila* | *warscewiczii* | Arecaceae |
| 40 | Cucsp | *Cucurbitaceae* | *sp* | Cucurbitaceae |
| 41 | Cupgla | *Cupania* | *glabra* | Sapindaceae |
| 42 | Cupliv | *Cupania* | *livida* | Sapindaceae |
| 43 | Davkun | *Davila* | *kunthii* | Dilleniaceae |
| 44 | Davnit | *Davila* | *nitida* | Dilleniaceae |
| 45 | Denarb | *Dendropanax* | *arboreus* | Araliaceae |
| 46 | Dicamp | *Dicranostyles* | *ampla* | Convolvulaceae |
| 47 | Dilsp | *Dillacarapacea* | *sp* | Dillacarapaceae |

| | | | | |
|---|---|---|---|---|
| 48 | Dolmul | *Doliocarpus* | *multiflorus* | Dilleniaceae |
| 49 | Dugsp | *Duguetia* | *sp* | Annonaceae |
| 50 | Eugsar | *Eugenia* | *sarapiquensis* | Myrtaceae |
| 51 | Eugsp | *Eugenia* | *sp* | Myrtaceae |
| 52 | Eutole | *Euterpe* | *oleracea* | Arecaceae |
| 53 | Eutpre | *Euterpe* | *precatoria* | Arecaceae |
| 54 | Fargla | *Faramea* | *glandulosa* | Rubiaceae |
| 55 | Farsp | *Faramea* | *sp* | Rubiaceae |
| 56 | Frisch | *Fridericia* | *schumanniana* | Bignoniaceae |
| 57 | Galsp | *Gallesia* | *sp* | Lauraceae |
| 58 | Geocon | *Geonoma* | *congesta* | Arecaceae |
| 59 | Goemei | *Goethalsia* | *meiantha* | Malvaceae |
| 60 | Goulup | *Gouania* | *lupuloides* | Rhamnaceae |
| 61 | Goupol | *Gouania* | *polygama* | Rhamnaceae |
| 62 | Guaaer | *Guatteria* | *aeruginosa* | Annonaceae |
| 63 | Guaamp | *Guatteria* | *amplifolia* | Annonaceae |
| 64 | Guadio | *Guatteria* | *diospyroides* | Annonaceae |
| 65 | Guagui | *Guarea* | *guidonia* | Meliaceae |
| 66 | Guarec | *Guatteria* | *recurvisepala* | Annonaceae |
| 67 | Hamapp | *Hampea* | *appendiculata* | Malvaceae |
| 68 | Heisca | *Heisteria* | *scandens* | Olacaceae |
| 69 | Helapp | *Heliocarpus* | *appendiculatus* | Malvaceae |
| 70 | Herdid | *Hernandia* | *didymantha* | Hernandiaceae |
| 71 | Hetsp | *Heteropteris* | *sp* | Malpighiaceae |
| 72 | Ilesku | *Ilex* | *skutchii* | Aquifoliaceae |
| 73 | Ingalb | *Inga* | *alba* | Leguminosae |
| 74 | Ingexa | *Inga* | *exalata* | Leguminosae |
| 75 | Ingoer | *Inga* | *oerstediana* | Leguminosae |
| 76 | Ingpun | *Inga* | *punctata* | Leguminosae |
| 77 | Ingsp | *Inga* | *sp* | Leguminosae |
| 78 | Ingthi | *Inga* | *thibaudiana* | Leguminosae |
| 79 | Ingumb | *Inga* | *umbellifera* | Leguminosae |
| 80 | Iridel | *Iriartea* | *deltoidea* | Arecaceae |
| 81 | Jaccop | *Jacaranda* | *copaia* | Bignoniaceae |
| 82 | Jubwil | *Jubelina* | *wilburii* | Malpighiaceae |
| 83 | Laepro | *Laetia* | *procera* | Salicaceae |
| 84 | Licmis | *Licaria* | *misantlae* | Lauraceae |
| 85 | Lozpit | *Lozania* | *pittieri* | Lacistemataceae |
| 86 | Mabocc | *Mabea* | *occidentalis* | Euphorbiaceae |
| 87 | Macsen | *Machaerium* | *senmani* | Leguminosae |
| 88 | Manhir | *Mandevilla* | *hirsuta* | Apocynaceae |
| 89 | Marnic | *Maripa* | *nicaraguensis* | Convolvulaceae |
| 90 | Mensp | *Mendoncia* | *sp* | Acanthaceae |
| 91 | Necmem | *Nectandra* | *membranacea* | Lauraceae |
| 92 | Ococer | *Ocotea* | *cernua* | Lauraceae |
| 93 | Ormsub | *Ormosia* | *subsimplex* | Leguminosae |
| 94 | Paslob | *Passiflora* | *lobata* | Passifloraceae |
| 95 | Paugra | *Paullinia* | *granatensis* | Sapindaceae |
| 96 | Paugrn | *Paullinia* | *grandifolia* | Sapindaceae |
| 97 | Pauing | *Paullinia* | *ingifolia* | Sapindaceae |
| 98 | Pauobo | *Paullinia* | *obovata* | Sapindaceae |

| | | | | |
|---|---|---|---|---|
| 99 | Pausp | *Paullinia* | *sp* | Sapindaceae |
| 100 | Penmac | *Pentaclethra* | *macroloba* | Leguminosae |
| 101 | Phagui | *Phanera* | *guianensis* | Leguminosae |
| 102 | Phopul | *Pholidostachys* | *pulchra* | Arecaceae |
| 103 | Pincor | *Pinzona* | *coriacea* | Dilleniaceae |
| 104 | Pippoe | *Piptocarpha* | *poeppigiana* | Compositae |
| 105 | Plusti | *Plukenetia* | *stipellata* | Euphorbiaceae |
| 106 | Posgra | *Posoqueria* | *grandiflora* | Rubiaceae |
| 107 | Poubic | *Pourouma* | *bicolor* | Urticaceae |
| 108 | Poudur | *Pouteria* | *durlandii* | Sapotaceae |
| 109 | Poumin | *Pourouma* | *minor* | Urticaceae |
| 110 | Procos | *Protium* | *costaricense* | Burseraceae |
| 111 | Propan | *Protium* | *panamense* | Burseraceae |
| 112 | Propit | *Protium* | *pittieri* | Burseraceae |
| 113 | Prorav | *Protium* | *ravenii* | Burseraceae |
| 114 | Psybra | *Psychotria* | *brachiata* | Rubiaceae |
| 115 | Psyela | *Psychotria* | *elata* | Rubiaceae |
| 116 | Psymar | *Psychotria* | *marginata* | Rubiaceae |
| 117 | Psyoff | *Psychotria* | *officinalis* | Rubiaceae |
| 118 | Psypan | *Psychotria* | *panamensis* | Rubiaceae |
| 119 | Psyrac | *Psychotria* | *racemosa* | Rubiaceae |
| 120 | Psysue | *Psychotria* | *suerrensis* | Rubiaceae |
| 121 | Pteroh | *Pterocarpus* | *rohrii* | Leguminosae |
| 122 | Quaoch | *Quararibea* | *ochrocalyx* | Malvaceae |
| 123 | Ransp | *Randia* | *sp* | Rubiaceae |
| 124 | Reisp | *Reinhardtia* | *sp* | Arecaceae |
| 125 | Rhokun | *Rhodostemonodaphne* | *kunthiana* | Lauraceae |
| 126 | Rhyery | *Rhynchosia* | *erythrinoides* | Leguminosae |
| 127 | Ricdre | *Richeria* | *dressleri* | Phyllanthaceae |
| 128 | Rindef | *Rinorea* | *deflexiflora* | Violaceae |
| 129 | Rolpit | *Rollinia* | *pittieri* | Annonaceae |
| 130 | Rolsp | *Rollinia* | *sp* | Annonaceae |
| 131 | Ryaspe | *Ryania* | *speciosa* | Salicaceae |
| 132 | Sensp | *Senegalia* | *sp* | Leguminosae |
| 133 | Sergon | *Serjania* | *goniocarpa* | Sapindaceae |
| 134 | Serpyr | *Serjania* | *pyramidata* | Sapindaceae |
| 135 | Sersp | *Serjania* | *sp* | Sapindaceae |
| 136 | Simama | *Simarouba* | *amara* | Simaroubaceae |
| 137 | Sipthe | *Siparuna* | *thecaphora* | Siparunaceae |
| 138 | Smidom | *Smilax* | *domingensis* | Smilacaceae |
| 139 | Smimol | *Smilax* | *mollis* | Smilacaceae |
| 140 | Smipur | *Smilax* | *purhampuy* | Smilacaceae |
| 141 | Smisp | *Smilax* | *sp* | Smilacaceae |
| 142 | Socexo | *Socratea* | *exorrhiza* | Arecaceae |
| 143 | Solsp | *Solanum* | *sp* | Solanaceae |
| 144 | Spasp | *Spachea* | *sp* | Malpighiaceae |
| 145 | Stesp | *Stemmadenia* | *sp* | Apocynaceae |
| 146 | Strmic | *Stryphodendron* | *microstachyum* | Leguminosae |
| 147 | Stylin | *Stygmaphyllum* | *lindenianum* | Malpighiaceae |
| 148 | Swacos | *Swartzia* | *costaricensis* | Leguminosae |
| 149 | Taccos | *Tachigali* | *costaricensis* | Leguminosae |

| | | | | |
|---|---|---|---|---|
| 150 | Tanpyr | *Tanaecium* | *pyramidatum* | Bignoniaceae |
| 151 | Tapgui | *Tapirira* | *guianensis* | Anacardiaceae |
| 152 | Terama | *Terminalia* | *amazonia* | Combretaceae |
| 153 | Tethyd | *Tetracera* | *hydrophila* | Dilleniaceae |
| 154 | Tetpan | *Tetragastris* | *panamensis* | Burseraceae |
| 155 | Thitom | *Thinouia* | *tomocarpa* | Sapindaceae |
| 156 | Triles | *Trichospermum* | *lessertianum* | Malvaceae |
| 157 | Trimex | *Trichospermum* | *mexicanum* | Malvaceae |
| 158 | Trisep | *Trichilia* | *septentrionalis* | Meliaceae |
| 159 | Troinv | *Trophis* | *involucrata* | Moraceae |
| 160 | Trorac | *Trophis* | *racemosa* | Moraceae |
| 161 | Unopit | *Unonopsis* | *pittieri* | Annonaceae |
| 162 | Unosp | *Unonopsis* | *sp* | Annonaceae |
| 163 | Virkos | *Virola* | *koschnyi* | Myristicaceae |
| 164 | Virseb | *Virola* | *sebifera* | Myristicaceae |
| 165 | Virsp | *Virola* | *sp* | Myristicaceae |
| 166 | Visbac | *Vismia* | *baccifera* | Hypericaceae |
| 167 | Visbil | *Vismia* | *billbergiana* | Hypericaceae |
| 168 | Vitcoo | *Vitex* | *cooperi* | Lamiaceae |
| 169 | Vocfer | *Vochysia* | *ferruginea* | Vochysiaceae |
| 170 | Vocgua | *Vochysia* | *guatemalensis* | Vochysiaceae |
| 171 | Vouano | *Vouarana* | *anomala* | Sapindaceae |
| 172 | Welreg | *Welfia* | *regia* | Arecaceae |
| 173 | Xylboc | *Xylopia* | *bocatorena* | Annonaceae |
| 174 | Xylser | *Xylopia* | *sericea* | Annonaceae |
| 175 | Xylsei | *Xylopia* | *sericophylla* | Annonaceae |
| 176 | Xylsp | *Xylosma* | *sp* | Salicaceae |
| 177 | Zanekm | *Zanthoxylum* | *ekmanii* | Rutaceae |
| 178 | Zansp | *Zanthoxylum* | *sp* | Rutaceae |

**Table S2.** Comparisons of species composition of the seed rain across four successional and mature forest plots over two time periods in Sarapiquí, Costa Rica. Summary statistics are for post-hoc tests after a significant pMANOVA test between successional and mature forests at each time period (1997-1999 and 2015-2017), between successional forests at each of the two time periods, and within successional forests across the two time periods, based on Holm-Bonferroni pairwise comparisons. Successional forests plots were sampled over two time periods; time period 1, in 1997-1999 (A1-D1 forest plots) and time period 2 in 2015-2017 (A2-D2 forest plots), and the mature forest (M) was sampled only in 2015-2017. All the pairs of forest plots statistically different from each other, after adjustment for multiple comparisons, are in bold under the *p adj* column.

| **Successional vs. Mature Forest in 1997-1999** | | | |
|---|---|---|---|
| **Plots** | ***F*** | ***p*** | ***p adj*** |
| A1 and M | 10.27 | 0.001 | **0.036** |
| B1 and M | 11.68 | 0.001 | **0.036** |
| C1 and M | 10.98 | 0.001 | **0.036** |
| D1 and M | 10.55 | 0.001 | **0.036** |
| **Successional vs. Mature Forest in 2015-2017** | | | |
| **Plots** | ***F*** | ***p*** | ***p adj*** |
| B2 and M | 3.08 | 0.002 | **0.036** |
| C2 and M | 2.75 | 0.009 | **0.039** |
| D2 and M | 2.57 | 0.019 | **0.041** |
| A2 and M | 9.64 | 0.001 | **0.036** |
| **Across successional forests in 1997-1999** | | | |
| **Plots** | ***F*** | ***p*** | ***p adj*** |
| A1 and B1 | 5.56 | 0.002 | **0.039** |
| B1 and C1 | 5.28 | 0.001 | **0.036** |
| B1 and D1 | 3.96 | 0.002 | **0.038** |
| C1 and D1 | 4.12 | 0.001 | **0.036** |
| **Across successional forests in 2015-2017** | | | |
| **Plots** | ***F*** | ***p*** | ***p adj*** |
| A2 and B2 | 7.54 | 0.001 | **0.036** |

| | | | |
|---|---|---|---|
| B2 and C2 | 2.86 | 0.004 | **0.037** |
| B2 and D2 | 2.18 | 0.003 | **0.036** |
| C2 and D2 | 2.18 | 0.052 | 0.067 |
| **Within successional forests between 1997-1999 vs. 2015-2017** | | | |
| **Plot** | ***F*** | ***p*** | ***p adj*** |
| A1 and A2 | 9.88 | 0.001 | **0.036** |
| B1 and B2 | 9.05 | 0.001 | **0.036** |
| C1 and C2 | 3.24 | 0.024 | **0.044** |
| D1 and D2 | 6.86 | 0.001 | **0.036** |

**Table S3.** Percentage and number of observed and shared species present in the seed rain in forests of different successional ages compared with mature forest in Sarapiquí, Costa Rica. For forest successional age, A1, B1, C1, D1 and A2, B2, C2, D2 represent the 1997-1999 and 2015-2017 time periods, respectively. Values inside parenthesis reflect the age of the successional forest at 1997 and 2017. *Total no. species* is the number of species present in the seed rain. *Unique species No. (%)* is the number and percentage of species respectively that are unique in the seed rain. *All species No. (%)* is the total number and percentage of species respectively which are present in the seed rain. *Shared species No. (%)* is the total number and percentage of species respectively which are present in both the successional and the mature forest. All percentages are calculated based on the total number of species.

| **Forest successional age** | **Total no. species** | **Successional forests** | **Mature Forest** | | **Shared species No. (%)** |
|---|---|---|---|---|---|
| | | **Unique species No. (%)** | **All species No. (%)** | **Unique species No. (%)** | |
| A1 (12) | 86 | 35 (41%) | 51 (59%) | 35 (41%) | 16 (18%) |
| B1 (15) | 90 | 39 (43%) | 51 (57%) | 42 (47%) | 9 (10%) |
| C1 (20) | 88 | 37 (42%) | 51 (58%) | 36 (41%) | 15 (17%) |
| D1 (25) | 87 | 36 (41%) | 51 (59%) | 38 (44%) | 13 (15%) |
| A2 (32) | 70 | 19 (27%) | 51 (73%) | 23 (33%) | 28 (40%) |
| B2 (35) | 71 | 20 (28%) | 51 (72%) | 39 (55%) | 12 (17%) |
| C2 (40) | 69 | 18 (26%) | 51 (74%) | 31 (45%) | 20 (29%) |
| D2 (45) | 67 | 16 (24%) | 51 (76%) | 34 (51%) | 17 (25%) |

**Figure S1.** Map of the study region in Sarapiquí, Costa Rica, showing the locations of the five forest plots of different successional ages (A-D; red dots) and the mature forest (M; blue dot). Successional plots were sampled in two time periods, 1997-1999 with plot ages varying from 12 to 25 in 1997 and 2015-2017 with plots ages varying from 32 to 45 years old in 2017. The mature forest was sampled only in 2015-2017. See Table 1 in the main text for the details for each forest plot. The map was constructed using the *get_map* function in the ggmap package (Kahle and Wickham 2013).

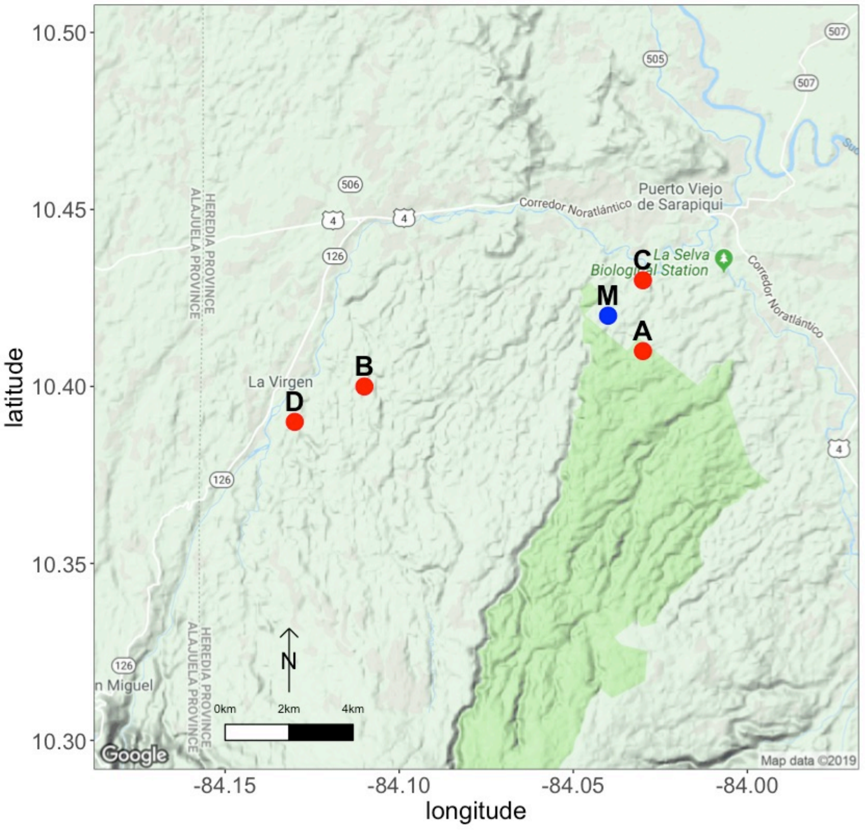

**Figure S2.** (a) Variation in the seed rain across four successional forest plots by months over two time periods in Sarapiquí, Costa Rica. Successional plots were sampled in two time periods, 1997-1999 (A-D) with plot ages varying from 12 to 25 in 1997 and 2015-2017 (A-D) with plots ages varying from 32 to 45 years old in 2017 (Table 1). Different colors represent the different forest plots; the 1997-1999 and 2015-2017 data are represented by a continuous and a dotted line, respectively. (b) Differences in average abundance of seeds rain across four successional forest plots and at two time periods (1997 – 1999 in red and 2015 – 2017 in blue) in Sarapiquí, Costa Rica, after controlling for variation in seed abundance by species. See Table 1 in the main text for details about each forest plot.

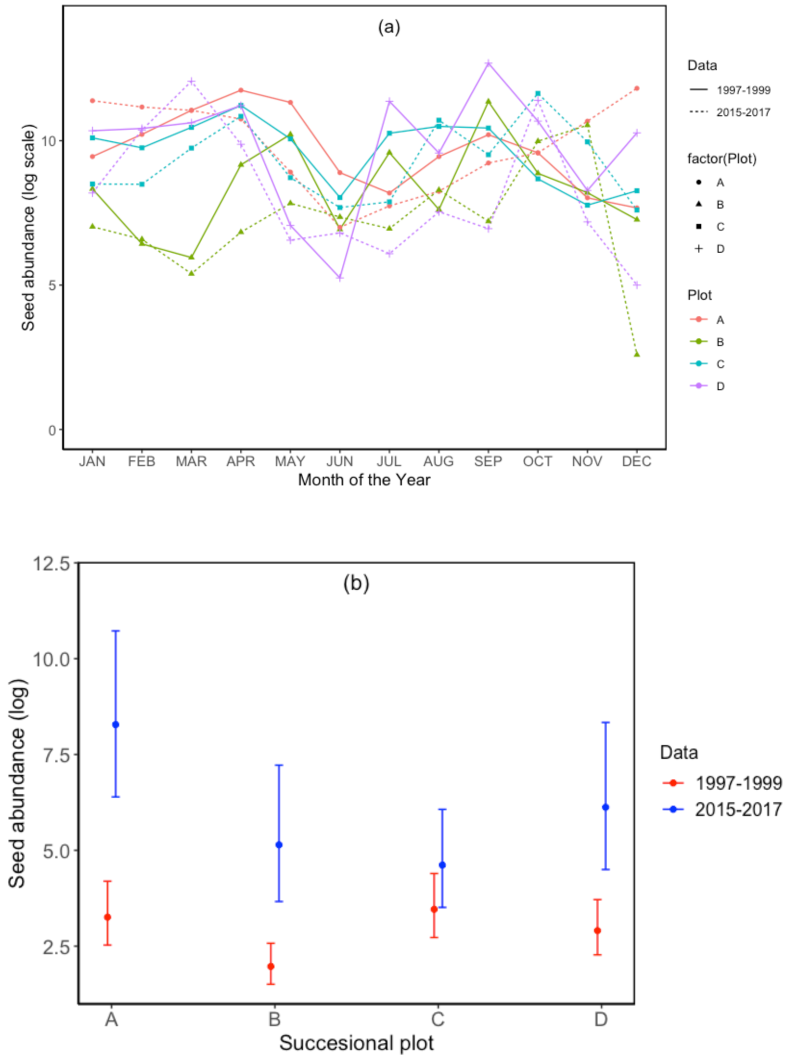

**Figure S3.** Abundance of woody species in the seed rain summed across four secondary forest plots in Sarapiquí, Costa Rica. The figure shows the 32 species that are higher or lower than the average seed abundance across all species.  Nine species were significantly lower in abundance, and 23 species were higher in abundance, than the mean abundance across all species. Species abbreviations correspond to the first three letters of the genus and species (Table S1).

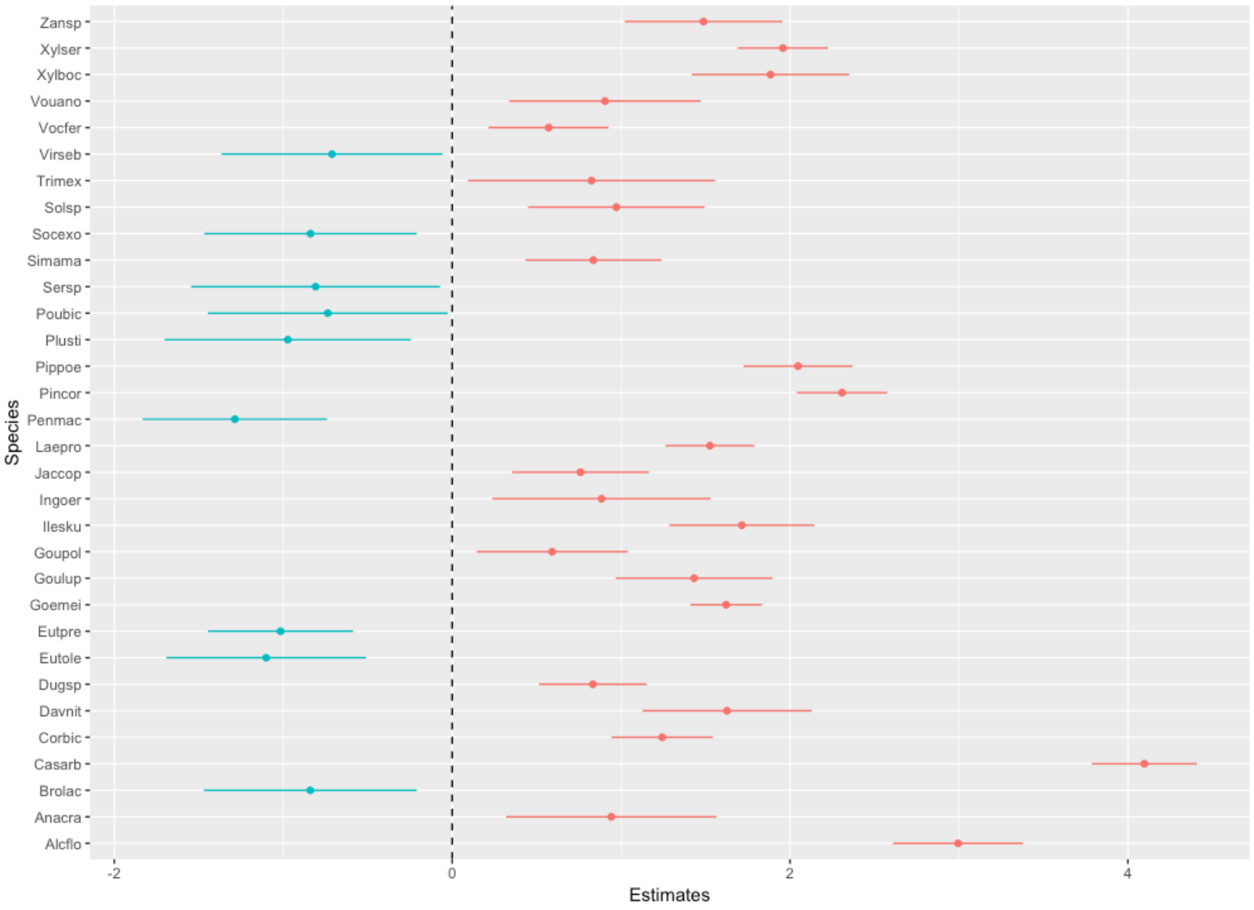

**Figure S4.** Abundance of seed rain across all secondary forests plots for species represented in seed traps continuously during all twelve months during 1997-1999 (top panel) and 2015-2017 (bottom panel) in Sarapiquí, Costa Rica. Species in 1997-1999 are light-demanding lianas (*Aristolochia sprucei* and *Pinzona coriacea*) or light-demanding tree species (*Cordia bicolor* and *Zanthoxylum sp*). Species in 2015-2017 are two shade tolerant palms (*Euterpe precatoria* and *Welfia regia*) and two light-demanding trees (*Goethalsia meiantha* and *Laetia procera*). Species abbreviations correspond to the first three letters of the genus and species (Table S1). See Table 1 in the main text for details about the successional forest plots.

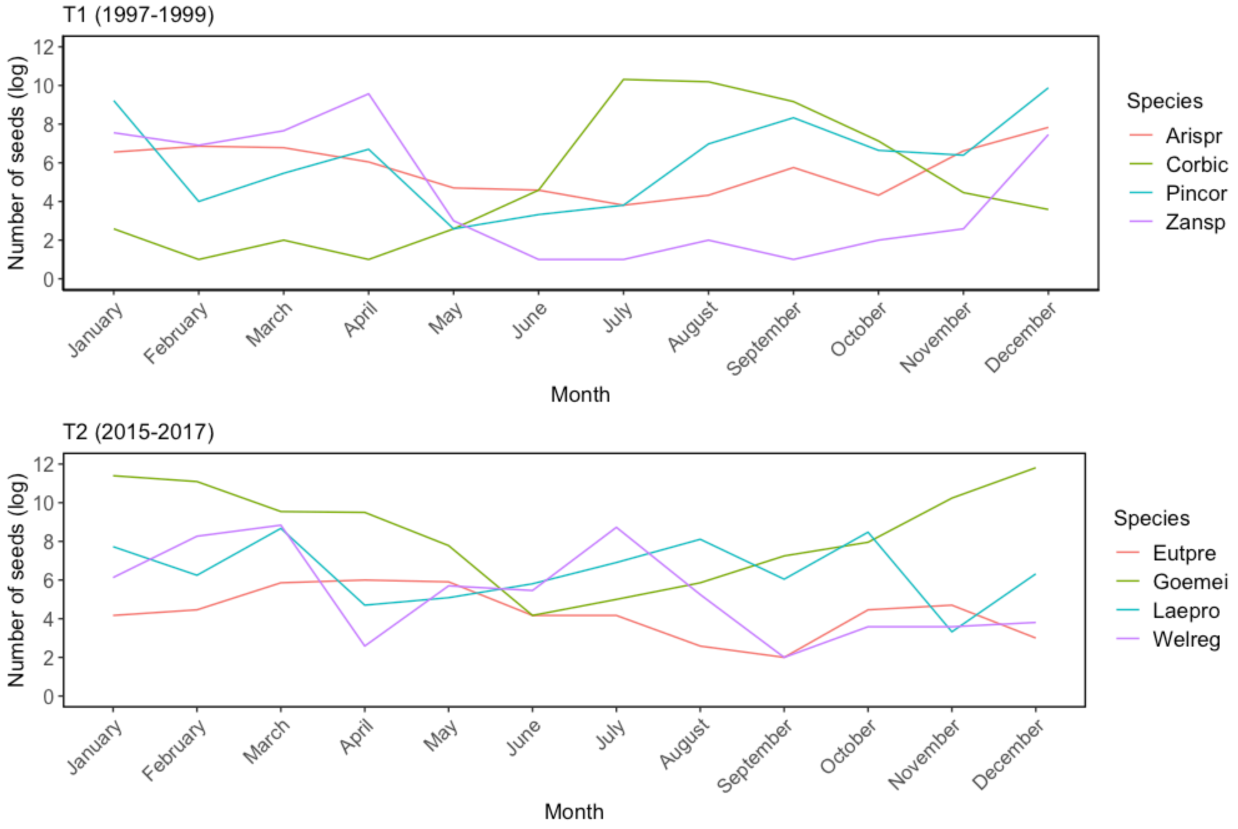

**Figure S5.** Null distributions of expected random Schoener's (*Cij*) co-occurrence indexes for species in successional vs mature forest plots, compared to the observed value, for each plot of different successional age. A1, B1, C1, D1 and A2, B2, C2, D2 represent successional plots in the 1997-1999 and 2015-2017 time periods, respectively. The red line represents the observed value, and the blue line is the mean, and the dotted blue lines are the 2.5th and 97.5th quantiles, of the null distribution. The standardized effect size of the co-occurrence index was calculated as the deviation of the observed value from the mean of the null distribution divided by the standard deviation of the null distribution.

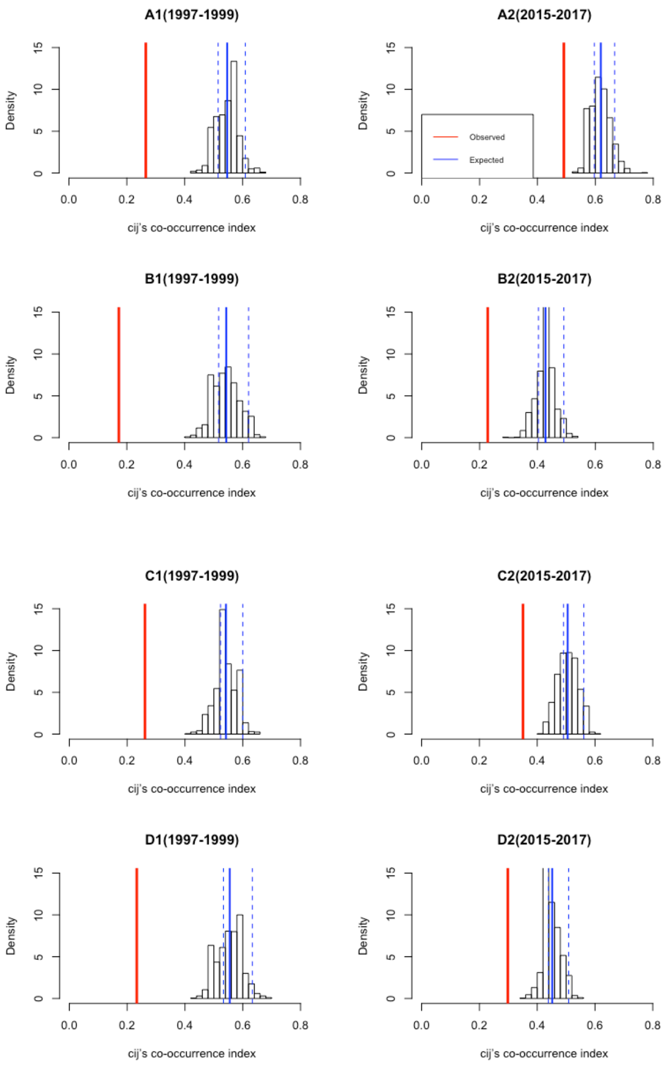

**Figure S6.** Species accumulation curves based on rarefaction of the seed rain in four successional (A-D) and mature (M) forest plots in Sarapiquí, Costa Rica. Successional plots were sampled in 1997-1999 (A1-D1 in red) and 2015-2017 (A2-D2 in blue), and the mature forest (M in black) was sampled only in 2015-2017. Plot successional ages: A1 = 12 B1 = 15, C1 = 20, D1 = 25, A2 = 32, B2 = 35, C2 = 40, and D2 = 45 years old. (a) small seeds (≤ 6 mm) and (b) large seeds (> 6 mm); (c) seeds of light demanding and (d) species shade tolerant species; (e) seeds of non-animal dispersed and (f) animal-dispersed species.

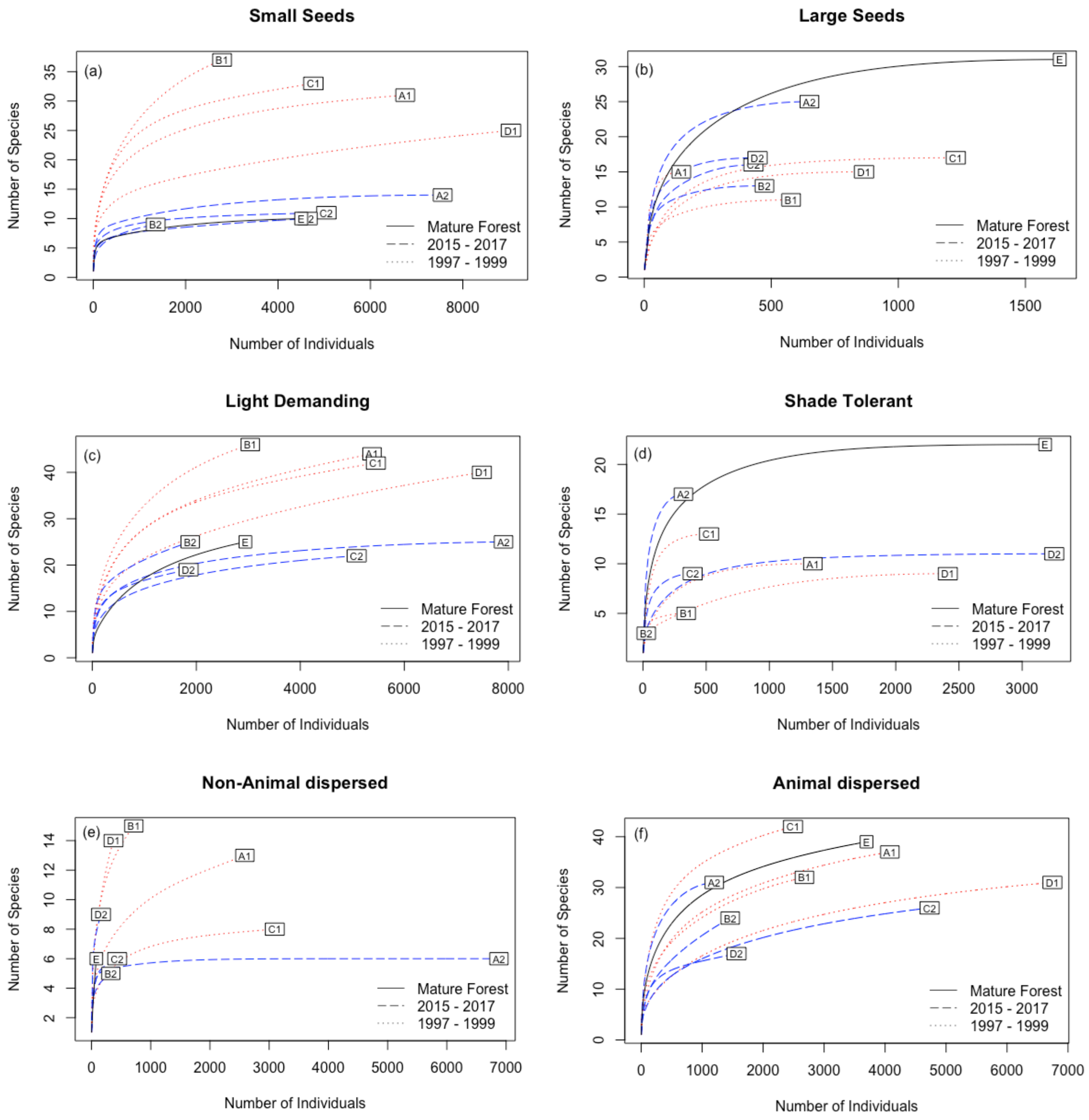

**Figure S7.** Decreasing proportion of immigrant seeds in secondary forests of increasing successional age in Sarapiquí, Costa Rica. The black line is the best fit line from ordinary least squares regression, and the gray shading indicates the confidence interval on the fit.

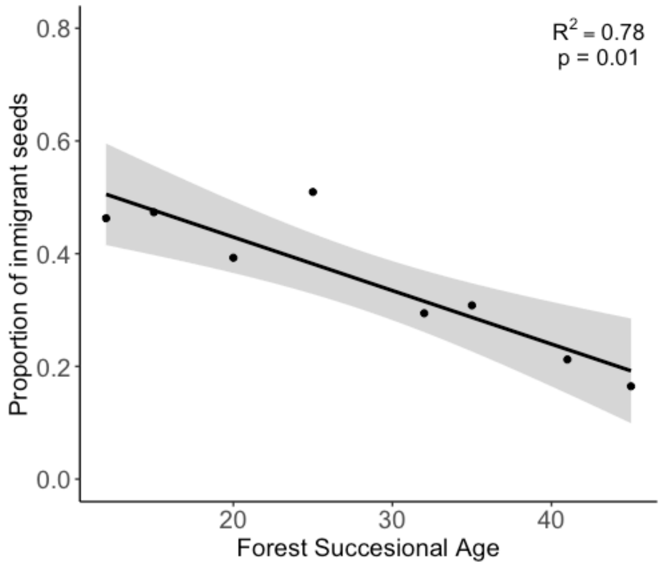
